# SARS-CoV-2 Sublingual Vaccine with RBD Antigen and Poly(I:C) Adjuvant: Preclinical Study in Cynomolgus Macaques

**DOI:** 10.1101/2022.09.21.508816

**Authors:** Tetsuro Yamamoto, Masanori Tanji, Fusako Mitsunaga, Shin Nakamura

## Abstract

Mucosal vaccine for sublingual route was prepared with recombinant SARS-CoV-2 spike protein receptor binding domain (RBD) antigen and poly(I:C) adjuvant components. The efficacy of this sublingual vaccine was examined using Cynomolgus macaques. Nine of the macaque monkeys were divided into three groups of three animals; control (just 400 μg poly(I:C) per head); low dose (30 μg RBD and 400 μg poly(I:C) per head); and high dose (150 μg RBD and 400 μg poly(I:C) per head), respectively. N-acetylcysteine (NAC), a mild reducing agent losing mucin barrier, was used to enhance vaccine delivery to mucosal immune cells. RBD-specific IgA antibody secreted in pituita was detected in two of three monkeys of the high dose group and one of three animals of the low dose group. RBD-specific IgG and/or IgA antibodies in plasma were also detected in these monkeys. These indicated that the sublingual vaccine stimulated mucosal immune response to produce antigen-specific secretory IgA antibodies in pituita and/or saliva. This sublingual vaccine also affected systemic immune response to produce IgG (IgA) in plasma. Little RBD-specific IgE was detected in plasma, suggesting no allergic antigenicity of this sublingual vaccine. Thus, SARS-CoV-2 sublingual vaccine consisting of poly(I:C) adjuvant showed reasonable efficacy in a non-human primate model.

## 1. Introduction

At the inception of the COVID-19 crisis, gene-based vaccine platforms, such as mRNA and DNA, brought a speed advantage. These gene-based vaccines were inserted a fragment of genetic code, which the cells must read to synthesize the proteins for themselves, along with the expression vector [1]. This has been preferred to rare but potentially vice-reactions, such as fever, headache, nausea, or chills. As protein-based vaccines have more good points, they are used to protect against hepatitis and other viral infections [2]. Although it needs much time to establish the protein vaccine to SARS-CoV-2, the protein-based vaccine is expected to become a mainstay in protecting the world from COVID-19, finally [3, 4].

To elicit a protective immune response by the protein-based vaccine, an immunity-stimulating adjuvant is indispensable along with protein antigen. Although there are several adjuvants, two are characteristics. One, MF59 or AS03, is an oil-in-water nano-emulsion stimulating Th1/Th2 [5]. The other is a double strand (ds)RNA poly(I:C), which is ligand for Toll-like receptor (TLR) 3 to activate immune and proinflammatory responses [6]. MF59 and AS03 were approved as adjuvants serving as intramuscular injected vaccine for influenza. Poly(I:C) is not yet approved due to its side effects of fever and proinflammatory cytokine production.

In addition to effective adjuvant, vaccination route is also a limited factor to establish protein-based SARS-CoV-2 vaccine. Since the coronavirus, like influenza, infects bronchial and alveolar epithelial cells, it is important to induce the secretion of virus antigen-specific IgA in the mucosa rather than IgG in the blood [7]. Recently, vaccines administered via alternative routes, such as nasal or oral, have been developed to elicit mucosal immune responses that differ from the systemic one [8]. Vaccinations through these routes often show higher efficacy than conventional subcutaneous vaccinations. Although nasal vaccines have been established and partly employed for clinical use [9], unpreferable influences to brain/central nerve system or lung were reported by its nasal administration [10-12]. On the one hand oral/sublingual vaccine revealed reasonable efficacy and high safety without the influences to brain [13]. In primates, humans and monkeys, the sublingual region has structural characteristic of wide space and is easily acceptable for vaccination rather than nasal space. These are advantageous reasons to choose sublingual roots for mucosal immune response. Furthermore, the above-mentioned side effects of poly(I:C) adjuvant were reported in nasal vaccination using rodent model [11,12,14]. These side effects would be affected with differences of adjuvant reactivity between rodents and primates due to the dissimilarity in their immune and related systems [15]. Different vaccine roots, nasal and sublingual, is also considered to influence the side effects.

This study examined sublingual vaccination using SARS-CoV-2 RBD antigen and poly(I:C) adjuvant in Cynomolgus monkeys. The objective of this study was to assess the efficacy of the poly(I:C) adjuvant in our sublingual conditions, in which NAC was used to disintegrate mucin barrier. In two monkey groups that were given low and high RBD antigen doses, RBD-specific IgA and IgG antibodies were detected in their pituita and plasma, respectively. These provided positive results for further study on the safety and efficacy of sublingual vaccines to SARS-CoV-2 using the monkey model.

## 2. Materials and Methods

### 2.1. Reagents and antibodies

N-acetylcysteine (NAC), bovine serum albumin (BSA), Na-Casein, sodium azido (NaN_3_), and Tween 20 are products of Fuji film-Wako (Japan). Phosphate-buffered saline (PBS; Nissui, Japan), Polyester swab (Nipponn Menbou, Japan), Filter spin column (Notgen Biotech, Canadian), Nunc-immune module, F8 Maxisorp (Thermo Fisher Scientific, USA), Streptavidin-HRP Conjugate (SA-HRP; Invitrogen, USA), and Tetramethyl benzidine (TMB; Sigma-Aldrich, USA) were used. Poly(I:C) HMW vaccine grade (poly(I:C); InvivoGen, USA) and Recombinant SARS CoV-2 Spike Protein Receptor Binding Domain (RBD; Creative Diagnostics, USA), and ELAST ELISA Amplification System (PerkinElmer, USA) was also employed.

Biotin-labeled (BT) monkey IgA antibody (Mabtech, Sweden), BT monkey IgA(alpha-chain) antibody (Merck, FRG), HRP-human IgG antibody (EY Laboratories, USA), and BT IgE antibody (Bio-Rad Laboratories, USA) were used.

### 2.2. Animals

Nine Cynomolgus macaque (Macaca fascicularis; male and female, 12.1 to 20.6 years old) were used here. Following the 3R policy of animal use, the macaque monkeys were reused by subsequential washout for 20 months after utilizing for subcutaneous injection of Sugi Basic Protein (Japanese Cedar Pollen Allergen). These monkeys were negative for B virus, SIV, TB, Shigella, Salmonella, and helminth parasites.

### 2.3. Vaccination and sampling

Poly(I:C) adjuvant(1 mg/ml)and RBD antigen (2 mg/ml) were kept at –70°C until use. Nine Cynomolgus macaques were divided into three groups of each three animals: control (mP01~03), low dose (mP04~06), and high dose (mP07~09). Each group’s animals were given the following vaccine formula, 0.7 ml of just 400 1g poly(I:C) per head for control;0.7 ml containing 30 μg RBD and 400 μg poly(I:C) per head for low dose;0.7 ml with 150 μg RBD and 400 μg poly(I:C) per head for high dose, respectively.

Before vaccination monkey’s sublingual surface was pretreated to disintegrate mucin layer for five minutes using wet cotton dipped in 1% NAC, and subsequently washed with saline. After wiping wet mucin surface with dry cotton, each 0.7 ml of vaccine material, control, low dose, or high dose, was administrated into sublingual space with a pipette and then allowed to stand for one minute at least.

These procedures for vaccination were conducted under anesthetization with mixture of medetomidine and ketamine and subsequent atipamezole to wake from the anesthesia. The sublingual vaccination was performed three times at four weeks interval. Sublingual booster was conducted 15 weeks after the 3rd vaccination to obtain samples for ELISA.

Blood and pituita were collected from each monkey under the above-mentioned anesthetization. Plasma samples were prepared after centrifugation of blood and used to assay RBD specific IgA, IgG, or IgE antibodies. Pituita samples adsorbed to a swab with polystyrene fiber were recovered by centrifugation using a spin-column and used for ELISA to measure RBD specific secretory IgA antibodies

### 2.4. ELISA

To detect RBD-specific IgA, IgG, or IgE antibodies, Nunc-immune module plates were coated with 100 μl of 5 μg/ml RBD in PBS by incubation at 37°C for one hr and then 4°C overnight. After washing with PBS-0.05% Tween 20, the plates were added with 1% Na-Casein in PBS-0.02% NaN_3_ for blocking, followed by incubation at 37°C for one hr and then kept at 4°C until use. Pituita or plasma samples were diluted 100 to 500-fold with 1% Na-casein-PBS-0.02% NaN_3._ These diluted samples were used as an ELISA sample. To perform ELISA, after removing the blocking reagent, the plates were added with each 50 μl of the diluted ELISA samples and 1M NaCl at a final concentration of 0.5 M to eliminate non-specific reaction.

After incubation at 37°C for one hr or at 4°C overnight and removing the samples, plates were washed with PBS-0.05% Tween 20. Then, detecting antibody, appropriately diluted BT-monkey IgA antibody, BT-monkey IgA (alpha-chain) antibody, HRP-human IgG antibody, or BT-IgE antibody was added, followed by incubation at 37°C for one hr. After washing, the plates were amplified using diluted SA-HRP and ELAST System mixture consisting of biotinyl tyramide. By this amplification, ELISA sensitivity was enhanced 10 to 30-fold at least.

After amplification plates were washed with PBS-0.05% Tween 20 and subsequently added with diluted SA-HRP, followed by incubation 37°C for one hr. Color development was performed with TMB and terminated by adding H_2_SO_4_, then absorption at 450 nm and 600 nm was read using a plate reader, iMark Microplate reader (Bio-Rad Laboratories, USA).

## 3. Results

In this study, two characteristic procedures were employed in ELISA and sublingual vaccine administration, respectively. One is the addition of NaCl at a final 0.5 M concentration into ELISA samples. This effectively eliminated non-specific biding events between the ELISA sample and RBD antigen, causing low background (data not shown). This non-specific event appeared to be an ionic interaction because of salt concentration dependency. The other was the prior treatment of the sublingual surface with NAC. NAC treatment loosens the mucinous layer that interferes with vaccine delivery to the sublingual immune system. NAC pretreatment yielded effective vaccination through mucin barrier reduction, as shown in Fig. 1.

**Figure 1.**
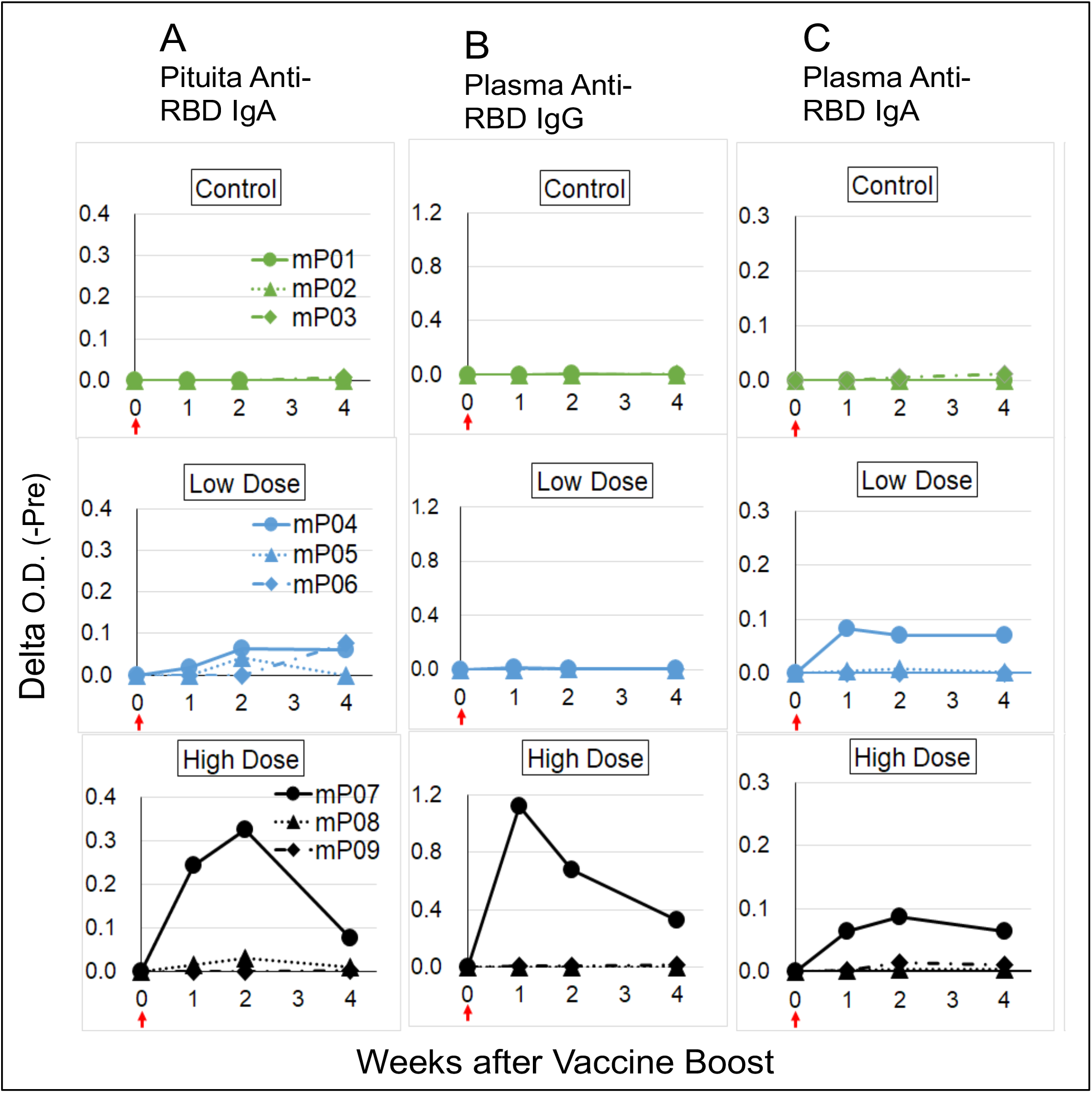
RBD-Specific antibodies induced by sublingual vaccination with SARS-CoV-2 RBD antigen and poly (I:C) adjuvant in Cynomolgus macaques. Three different vaccine doses of control (400 μg poly(I:C) per head), low dose (30 μg RBD and 400 μg poly(I:C) per head) and high dose (150 μg RBD and 400 μg poly(I:C) per head). A: RBD-specific IgA secreted in pituita, B; RBD-specific IgG in plasma, C; RBD-specific IgA in plasma. Red arrow; vaccine boost.

Sublingual SARS-CoV-2 vaccine consisting of RBD antigen and poly(I:C) adjuvant elicited local mucous and systemic immune response. Figure 1 shows the RBD-specific antibody titer by a boost administration following three times’ sublingual vaccination, IgA in pituita (A); IgG in plasma (B); and IgA in plasma (C). RBD-specific IgA was detected in pituita of both groups of low (30 μg/head) and high (150 μg/head) RBD dosage, indicating that the sublingual vaccine with poly(I:C) adjuvant induces antigen-specific secretory IgA production by mucous immune system in nose and/or mouth.

As seen in Fig. 1A (pituita anti-RBD IgA), the local mucous immune response, evoked by sublingual vaccination appeared to be in a dose-response manner. The responding monkeys’ numbers and their antibody titers differed between the low and high dose groups; one animal (mP04) with low titer in the low dose group and two animals (mP07 and mP08) with high and low titers in the high dose group. The dose-response was also observed in systemic immune response of plasma anti-RBD IgG, as shown in Fig. 1B, one animal (mP07) with high antibody titer in the high dose group but none in low dose groups. The plasma IgA antibody titer was poor in high (mP07) and low (mP04) dose animals.

No RBD-specific plasma IgE was detected (data not shown), suggesting that sublingual vaccine with poly(I:C) adjuvant had little vice-reactions to cause an allergic response. The sublingual vaccination neither raised flare and/or edema around the sublingual region nor decreased body weight and/or appetite.

## 4. Discussion

As mucosal vaccines offer the potential to trigger robust protective immune responses at the predominant sites of pathogen infection, practical vaccines against air-borne and/or droplet infectious viruses, such as SARS-CoV-2, should be administered into the nasal or oral cavity to establish an anti-virus mucus immunity to produce secretory IgA antibodies [16,17]. There is a thick mucinous layer comprised of soluble and fixed mucin on the inner surface of the nasal or oral cavity. This mucinous layer is a barrier that interferes with the interaction between vaccine material(s) and mucosal immune cells existing under the mucin [18]. Thus, effective vaccine delivery is an essential factor in developing mucosal vaccines.

The mucin layer comprises a highly O-glycosylated glycoprotein linked with disulfide binds [19]. NAC is a mild reducing reagent used as a drug, Mucofilin, for respiratory tract viscous liquid resolvent. NAC was also employed for mucin disintegration [20] and removal of nasal mucus [21]. NAC pretreatment yielded excellent results in previous examinations for bladder transplantation of cancer cells and nasal sensitization with cider pollen antigen using monkey model (data not shown). These are reasons why NAC for sublingual vaccination was employed. The different, ineffective result was reported in a previous study, in which sublingual vaccination was performed under similar conditions as those of use, monkey model and poly(I:C) adjuvant, except for non-use of NAC [22]. Another case of sublingual vaccination without mucin treatment was also reported to fail to induce specific IgA or IgG in rhesus monkeys [23]. Even though several factors yielded different results, a possible main factor would be the above-mentioned mucinous barrier.

The practical mucosal vaccine will be administered through the nasal or oral cavity route. In the case of nasal route, the vaccine is sprayed into the nasal cavity, where it is difficult to know the exact point and/or amount of administered vaccine. Conversely, the oral route, especially sublingual administration, is simple and convenient because of self-visualizing all of vaccination procedures, including site(s) and amount. Although many reports about nasal vaccine in rodents exists, little knowledge was accumulated on sublingual or oral one, especially in primates, except for three reports [22-24].

In this study, poly(I:C) is used for the sublingual vaccine because of its potent effects as a TLR3 ligand [6, 25]. Poly (I:C) is a dsRNA, consisting of a polyinosinic and polycytidylic acid. Its dsRNA nature mimics viral infection through binding endosomal TLR3 and cytosomal receptors retinoic acid-inducible gene I and melanoma differentiation-associated gene 5 [26, 27]. Poly(I:C) is known for its immunostimulatory activity due to its capacity to activate immune cell types [28]. Therefore, poly(I:C) is considered a potent vaccine adjuvant to activate antigen-presenting cells, particularly dendritic cells [29, 30]. Poly(I:C)-mediated TLR-3 activation leads to proinflammatory cytokines production and/or related factors, type I IFN, IL-15, NK [31]. Although in this context, clinical use of poly(I:C) as vaccine adjuvant has been unapproved yet except for limited cancer use, studies to develop its use for vaccine adjuvant is progressing in preclinical and clinical fields [32].

The poly(I:C)-mediated proinflammatory cytokines productions and related factors were mainly reported in studies using nasal vaccination in mice [11,12,14]. Differences in the immune system between rodents; mice and rat, and primates; humans and monkeys, were remarked by genome-based evidence [15]. As in mice, poly(I:C) is the most effective inducer of type I IFN among TLR agonists [33], its marked proinflammatory cytokine pathway activation might be over-estimated. It is also thought to be plausible that these poly(I:C)-mediated reactions differ at nasal and sublingual sites. Information on poly(I:C)-mediated vice-reactivity for its use in sublingual vaccine adjuvant is quite insufficient in case of primates, monkeys and humans, yet.

## 5. Conclusions

As part of developing a practical SARS-Cov-2 sublingual vaccine using poly(I:C) adjuvant, a preclinical study using the monkey model was performed. RBD-specific IgA antibody was detected in pituitas of both monkey groups given high (2 of 3) and low (1 of 3) antigen doses, respectively. From these, it was indicated that this sublingual vaccine could elicit mucosal immune response to produce secretory IgA antibodies to SARS-Cov-2. This study is yet to examine the exact safety and efficacy using genomic markers described in previous papers in mice [12]. Further studies on these points are in progress using the preclinical non-human primate model.

## Supplementary Information

### Author Contributions

Conceptualization, T. Y. and S.N.; methodology, F.M.; investigation, F.M and S.N.; resources, T. Y. and M.T.; data curation, F.M.; writing—original draft preparation, S.N. and F.M.; writing— review and editing, S.N. and T. Y.; visualization, F.M..; supervision and project administration, T. Y. and M.T.; funding, T. Y.; All authors have read and agreed to the published version of the manuscript.

### Funding

This study received no external funding.

### Institutional Review Board Statement

This study was conducted according to the guidelines of Institutional Animal Care and Committee Guide of Intelligence and Technology Lab, Inc. (ITL) based on the Guidelines for Proper Conduct of Animal Experiments and approved by the Animal Care Committee of the ITL (approved number: AE2021001, data: 7 July 2021). This study was also approved by the ITL Biosafety Committee (approved number: BS202100, date: 7 July 2021).

### Informed Consent Statement

Not applicable.

### Data Availability Statement

Data are available from S.N. upon reasonable request.

## Acknowledgments

We thank to Kazuhiro Kawai for his invaluable technical assistance.

## Preprint Information

After submitting this paper, we have posted information concerning safety and vice-reactivity/side effect of sublingual vaccine with poly (I:C) adjuvant on bioRxiv preprint [34]. We have also posted information concerning SARS-CoV-2 neutralization assay using plasma with anti-RBD IgA/IgG antibody on bioRxiv preprint [34].

---

**Supplemental Figure:**
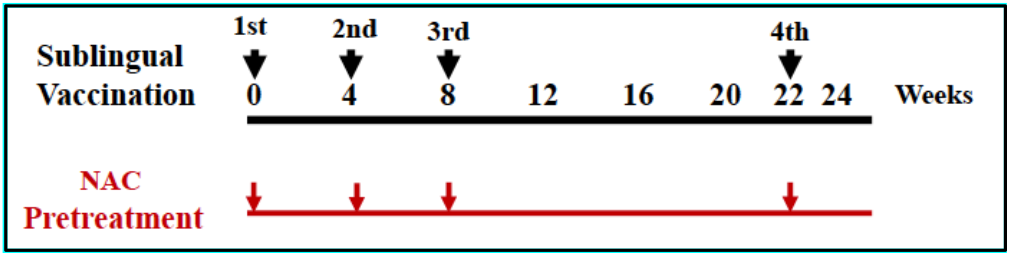
Outline protocols for sublingual vaccination and NAC pre-treatment in *Cynomolgus macaque*

## References

1. D. Pushparajah, et al. Advances in gene-based vaccine platforms to address the COVID-19 pandemic.Advanced Drug Delivery Reviews, 170 (2021) 113–141 .doi.org/10.1016/j.addr.2021.01.003

2. P.C. Soema, et al. Development of cross-protective influenza A vaccines based on cellular responses, Frontiers in Immunology, published: 15 May 2015, doi: 10.3389/fimmu.2015.00237.

3. Arashkia, et al. severe acute respiratory syndrome-coronavirus-2 spike (S) protein-based vaccine candidates: State of the art and future prospects, Rev Med Virol. 2020; e2183. doi.org/10.1002/rmv.2183.

4. Dolgin E., How protein-based COVID vaccines could change the pandemic. Nature. 2021 Nov;599(7885):359–360. doi: 10.1038/d41586-021-03025-0.

5. SC De Rosa, et al. Whole-blood cytokine secretion assay as a high-throughput alternative for assessing the cell-mediated immunity profile after two doses of an adjuvanted SARS-CoV-2 recombinant protein vaccine candidate, Clin Transl Immunology. 2022 Jan 11;11(1):e1360. doi: 10.1002/cti2.1360.eCollection 2022.

6. Hafner AM, et al. Particulate formulations for the delivery of poly(I:C) as vaccine adjuvant, Adv Drug Deliv Rev.2013 Oct;65(10):1386–99. doi: 10.1016/j.addr.2013.05.013.

7. Ainai, et al. Human immune responses elicited by an intranasal inactivated H5 influenza vaccine, Microbiology and Immunology. 2020; 64:313–325., DOI: 10.1111/1348-0421.12775 .

8. R. Mudgal, et al. Prospects for mucosal vaccine: shutting the door on SARS-CoV-2, Human Vaccine Immnunother., 2020, 16(12), 2921–2931, doi.org/10.1080/21645515.2020.1805992.

9. C.S Ambrose et al. Current status of live attenuated influenza vaccine in the United States for seasonal and pandemic influenza, Influenza and Other Respiratory Viruses, 2, 193–202 2008, doi: 10.1111/j.1750-2659.2008.00056.x.

10. Lemiale F, et al., Enhanced mucosal immunoglobulin A response of intranasal adenoviral vector human immunodeficiency virus vaccine and localization in the central nervous system, J.Virol.Sept. 2003, 10078–87 ; DOI: 10.1128/JVI.77.18.10078–10087.2003.

11. Sasaki E, et al. Establishment of a novel safety assessment method for vaccine adjuvant development, Vaccine, 36 (2018) 7112–7118, /doi.org/10.1016/j.vaccine.2018.10.009.

12. Sasaki E, et al., Immunogenicity and Toxicity of Different Adjuvants Can Be Characterized by Profiling Lung Biomarker Genes After Nasal Immunization, Frontiers in Immunolog, September 2020, doi: 10.3389/fimmu.2020.02171.

13. Song J-H, et al, Sublingual vaccination with influenza virus protects mice against lethal viral infection, Proc Natl Acad Sci. 2008 Feb 5;105(5):1644–9. doi: 10.1073/pnas.0708684105.

14. Sasaki E, et al. Modeling for influenza vaccines and adjuvants profile for safety prediction system using gene expression profiling and statistical tools, PLOS ONE, February 6, 2018, doi.org/10.1371/journal. Pone.0191896.

15. J. Mestas, C. C. W. Hughes, Of Mice and Not Men: Differences between Mouse and Human Immunology, J Immunol 2004; 172:2731–2738;doi: 10.4049/jimmunol.172.5.2731.

16. C Czerkinsky, J Holmgren, Mucosal delivery routes for optimal immunization: targeting immunity to the right tissues, Curr.Top Microbiol Immunol, 2012;354:1–18. doi: 10.1007/82_2010_112

17. Lavelle EC., Ward RW., Mucosal vaccines-fortifying the frontiers, Nat. Rev. Immunol (2021). 10.1038/41577-s021-00583-.2.

18. T.L. Carlson, J.Y. Lock, Engineering the Mucus Barrier, Annu Rev Biomed Eng. 2018 June 04; 20: 197–220. doi:10.1146/annurev-bioeng-062117-121156.

19. A.P. Corfield, Mucins: a biologically relevant glycan barrier in mucosal protection, Biochim Biophys Acta, 2015,1850(1):236–52. doi: 10.1016/j.bbagen.2014.05.003.

20. Pillai K., et al, A formulation for in situ lysis of mucin secreted in pseudomyxoma peritonei, Int. J. Cancer: 134, 478–486 (2014), doi: 10.1002/ijc.28380.

21. H. Suzuki, et al, Impaired airway mucociliary function reduces antigen-specific IgA immune response to immunization with a claudin-4-targeting nasal vaccine in mice, Sci. Rep. (2018) 8:2904 | DOI: 10.1038/s41598-018-21120-7.

22. Veazey RS., et al, Evaluation of mucosal adjuvants and immunization routes for the induction of systemic and mucosal humoral immune responses in macaques, Human Vaccines & Immunother., 11:12, 2913--2922; December 2015; doi.org/10.1080/ 2015.1070998

23. Alan D. Curtis II, et al, A simultaneous oral and intramuscular prime/sublingual boost with a DNA/Modified Vaccinia Ankara viral vector-based vaccine induces simian immunodeficiency virus-specific systemic and mucosal immune responses in juvenile rhesus macaques, J Med Primatol. 2018 October; 47(5): 288–297. D

24. Johnson S., *et.al*, 589. Oral Tablet Vaccination Induces Heightened Cross-Reactive CD8 T Cell Responses to SARS-COV-2 in Humans, Open Forum Infect Dis. 2021 Nov; 8 (Suppl 1): S397. doi: 10.1093/ofid/ofab466.787.

25. K. Tewari, *et.al*, Poly(I:C) is an effective adjuvant for antibody and multi-functional CD4+ T cell responses to Plasmodium falciparum circumsporozoite protein (CSP) and αDEC-CSP in Non-Human Primates, Vaccine. 2010 October 21; 28(45):7256–7266. doi: 10.1016/j.vaccine.2010.08.098.

26. Yu M, Levine SJ. Toll-like receptor, RIG-I-like receptors and the NLRP3 25. inflammasome: key modulators of innate immune responses to double-stranded RNA viruses. Cytokine Growth Factor Rev.2011;22(2):63–72. doi.org/10.1016/j.cytogfr.2011.02.001.

27. Waele, JD, et al, A systematic review on poly(I:C) and poly-ICLC in glioblastoma: adjuvants coordinating the unlocking of immunotherapy, J Exp Clin Cancer Res, 2021 Jun 25;40(1):213. doi: 10.1186/s13046-021-02017-2.

28. Ammi R, et al. Poly(I:C) as cancer vaccine adjuvant: knocking on the door of medical breakthroughs. Pharmacol Ther. 2015; 146:120–31. 10.1016/j.pharmthera.2014.09.010.

29. Fucikova J, et al. Poly I: C-activated dendritic cells that were generated in CellGro for use in cancer immunotherapy trials. J Transl Med. 2011;9(1):223. 10.1186/1479-5876-9-223.

30. Kato H, Takeuchi O, Sato S et al. (2006) Differential roles of MDA5 and RIG-I helicases in the recognition of RNA viruses. Nature. 441: 101–5. doi: 10.1038/nature04734.

31. Sultan H, et al, Role of MDA5 and interferon-I in dendritic cells for T cell expansion by anti-tumor peptide vaccines in mice, Cancer.Immunol. Immunother.,2018 Jul;67(7):1091–1103. doi: 10.1007/s00262-018-2164-6.

32. Naour JL, et al. Trial watch: TLR3 agonists in cancer therapy, Oncoimmunogy.2020, Jun 2;9(1):1771143. doi: 10.1080/2162402X.2020.1771143.

33. Longhi MP, et al. Dendritic cells require a systemic type I interferon response to mature and induce CD4+ Th1 immunity with poly IC as adjuvant. J Exp Med. 2009; 206(7):1589–602. 10.1084/jem.20090247.

34. Yamamoto T., et al. Mechanism Underlying the Immune Responses of a Sublingual Vaccine for SARS-CoV-2 with RBD Antigen and Adjuvant, Poly(I:C) or AddaS03, in Non-human Primates, bioRxiv preprint, 10.1101/2023.05.15.540684).

